# Impact of mitochondrial fission and fusion on neuronal health and incidence of dementia

**DOI:** 10.1101/2024.12.21.629890

**Authors:** Alan G Holt, Adrian M Davies

## Abstract

Mitochondria are highly dynamic structures that undergo constant remodeling by the process of fission and fusion. Using simulation methods we study the effects of neuron loss in humans due to the proliferation of deletion mutations. We implement two models of an organelle, namely, closed cristae (CC) and open mitochondrion (OM). With CC, mtDNA are confined to a cristae unless *mixed* by the process of fission and fusion. Conversely, mtDNA can diffuse freely throughout the organelle in the OM model. We also implement selective mitophagy in the CC model.

Higher rates of mixing mtDNA increase the rate of neuron loss which, in turn, increases the prevalence of dementia. Selective mitophagy mitigates against the effect of high rate mixing. However, this mitigation is all or nothing. Even at nominal rates compared to mixing, neuron loss is almost completely eliminated resulting in minimal cognitive decline. These results do not fit the observations of normal aging and dementia prevalence in the elderly human population.

This study brings into question the role that fission and fusion is believed play in the removal of defective mtDNA.

## 1 Introduction

Mitochondria continually undergo fission and fusion [8, 63]. Fission is the process by which a mitochondrion fragments into smaller mitochondria and fusion is where fragmented mitochondria merge into a single, larger mitochondrion. Conventional wisdom mandates that fission and fusion mixes mitochondrial contents [60, 2] throughout the mitochondrion, permitting complementation of mitochondrial DNA (mtDNA) and mtDNA products [43]. This helps to maintain mitochondrial function by distributing the energy-producing machinery [1, 45] enabling it to tolerate increasing levels of heteroplasmy and defective mtDNA [2]. Fission and fusion also facilitates cellular adaptation to changing energy demands and other metabolic needs [61]. It may also be necessary to remove damaged mitochondria through a process of selective mitophagy where dysfunctional mitochondrial fragments are expunged from the cell [61, 5].

In this paper we use simulation methods to investigate the long term effect fission and fusion has on the proliferation of deletion mutants (mtDNA_*del*_) within human neurons and the impact that it has on the prevalence of dementia in the elderly. We implement two separate mitochondrial models, namely, an open mitochondria model (OM) and a closed cristae model (CC) such that we can simulate them in software. In the OM model, mtDNA are free to randomly drift throughout the organelle but in the CC model mtDNA are confined to a particular cristae and can only move between cristae by means of fission and fusion. We run OM model simulations for comparison with CC.

For CC, we consider two features of fission and fusion and record their impact on neuron loss:

- Mixing *only*, 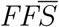. With 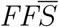, fissioned fragments from a cristae are always fused back to the main mitochondrion, albeit, to some other cristae chosen at random. The mtDNA in a fissioned fragment joins the mtDNA population of cristae it fuses with.
- Mixing with selective mitophagy, *FFS*. With *FFS*, fission fragments can only fuse back if they are sufficiently healthy. Unhealthy fragments undergo *selective mitophagy* and their mtDNA are expunged from the main mitochondrion.

We run simulations for various mixing rates 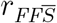 and selective mitophagy rates *r*_*FFS*_. The results of our simulation calls into question the roles that mixing and selective mitophagy play in mitochondrial dysfunction. We found for 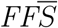, that cognitive decline increased with 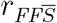, where high mixing rates brought forward the onset of dementia and increased severity. However, *FFS* eliminated neuron loss altogether such that hosts suffered no cognitive decline at all. Thus, our CC/*FFS* model vastly underestimated the prevalence of dementia observed in the population. Given how well selective mitophagy performed *in silico*, we are skeptical of its role *in vivo*. We tried a number of variants of *FFS* with a view to reducing the effectiveness of selective mitophagy, such as slowing down *r*_*FFS*_ (relative to 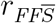) and significantly increasing a host’s predisposition to dementia (increasing deletion mutation rate). However, the resultant levels of heteroplasmy were far greater than empirical observations.

The contributions of this paper are three-fold:

- Simulates fission and fusion of mitochondria.
- Presents results of 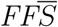 and *FFS* simulations and how neuron loss is impacted for various mixing and selective mitophagy rates.
- Calls into question the role that fission and fusion plays in the removal of defective mtDNA in mitochondria.

## 2 Background

Mitochondria can appear as a threadlike network or as distinct granules [7] with dynamic interconversion between these two forms through repeated fission and fusion [8, 14]. The architecture of the mitochondrion is as a set of cristae forming individual invaginations; compartmentalising mtDNA and their products [27]. The cristae provide the necessary surface area for the respiratory complexes [48], for localising the proton gradient [50] and (likely) maintaining the local concentration of metabolites [40].

In previous studies [25], we modeled the mitochondria as an open organelle (OM). In the simulation, each mtDNA in the population is processed sequentially. We removed any ordering bias by randomly sorting the mtDNA prior to each simulation cycle. Thus, in the OM model, mtDNA were considered free to disperse throughout the organelle, albeit, “mixing” was merely an artefact of the algorithm. In contrast, organelle-wide mixing in the CC model is entirely dependent upon the process of fission and fusion.

It is believed that fission and fusion enables even distribution of the mitochondrial content. Proteins coded by wild-type mtDNA (mtDNA_*wild*_) could, by complementation, serve to ameliorate the effects of defective or absent gene products from damaged mtDNA [61, 32, 39, 41, 8, 14].

It is widely accepted that mitochondria originated from ancestral bacterial species through endosymbiosis [37]. Given that the function of fusion in bacteria is to facilitate the exchange of genetic material [51], it is plausible that fission and fusion performs a similar function in eukaryotic cells.

In addition to mixing, fission and fusion is believed to act as a purification mechanism. Fusion of the mitochondrial fragment back to the main mitochondrion is dependent on the fragment being able to generate an adequate membrane potential (ΔΨ) [34, 38, 54]. Fragments with inadequate ΔΨ will fail to fuse and are tagged for mitophagy. This process of selective mitophagy would serve to maintain the mtDNA_*wild*_ population from being overwhelmed by mutant strains of mtDNA such as mtDNA_*del*_ [5, 11, 30, 20].

Thus, we consider two forms of fission and fusion within the CC model. With the first, a fissioned fragment *always* fuses back to the main mitochondrion 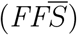. Albeit, to a different cristae from which it fissioned from, thereby, mixing mtDNA throughout the organelle. In the second form, the fragment only fuses back if it is deemed healthy (*FFS*).

### 2.1 Mixing

It is not clear how static or dynamic the cristae morphology is [29] but, for simulation purposes, we need some estimate of the rate of fission and fusion. Twig *et al*. [55] gives an average per-cell rate of 1 event every 15 minutes. Coincidentally, our simulation advances in intervals of *t* = 15 minutes, thus, 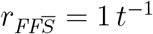, where 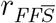 is the per-cell rate of fission and fusion. Mouli *et al*. [39] gives a similar per-cell value of *≈* 70 to 100 daily events (0.73 to 1.04 *t*^*−*1^). Cagalinec *et al*. [12] provides a slightly slower rate of 33 events per day (per cell). Jendrach *et al*. [28], however, gives a per-cell rate of 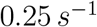 which would equate to 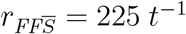.

The Jendrach [28] rate, at two orders of magnitude higher than those previously cited, appears to be an outlier here, thus, we run simulations over a range of 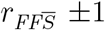 orders of magnitude of 1 *t*^*−*1^. Note, we also consider a rate 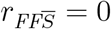 (no mixing).

In our (CC) model the fusion rate is always the same as the fission rate. We do not consider the condition whereby fission rate and fusion rate are different. Gene mutations that cause disparate rates in fission and fusion are, typically, associated with conditions such as Charcot-Marie-Tooth disease and motor neuropathy [16]. Other mutations in genes for fission and fusion can be fatal [6] but, unlike dementia, are not a function of age.

### 2.2 Selective Mitophagy

Selective mitophagy is a specialised form of autophagy [53], where damaged or dysfunctional parts of the mitochondria are specifically targeted and degraded by the cell. This process is believed to be crucial for maintaining cellular health and homeostasis by ensuring the removal of mitochondria that can no longer function properly [5]. In our simulation, we consider selective mitophagy to be integral to the fission and fusion process, and that a fragment may only fuse back if it can maintain a sufficient level of ΔΨ [34], if not, it is targeted for mitophagy.

Selective removal of damaged mtDNA, requires that, mtDNA must be localised with its gene products [33] in order to achieve a threshold effect [49] that is representative of the quality of the mtDNA population in a fragment. In the CC model, the components of the electron transport chain, and the mtDNA that encoded it, are assumed to be locally associated. Thus, when a fragment fissions from a cristae, the phenotype is a result of the genotype.

In contrast, the electron transport chain components for the OM model are not necessarily encoded by the local mtDNA at the time of a fission event and the phenotype is not necessarily indicative of the genotype. Thus, selective mitophagy necessitates a *closed* cristae environment in order to function, whereas an open mitochondrion would confound such a purification mechanism.

For most of our experiments, we consider the rate of selective mitophagy *r*_*FFS*_ to be the same as 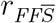. However, for the last experiment, we disassociate the two rates such that *r*_*FFS*_ is set much slower than 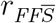.

## 3 Simulator

We simulate the proliferation of mtDNA_*del*_ amongst the mtDNA_*wild*_ species within a pseudo mitochondrial organelle. We describe the simulator in this section (written in Python) which is based upon the Mitochondrial Free Radical Theory of Aging (MFRTA). In our simulator mtDNA are software abstractions that, like their biological counterparts, are subject to *expiration, replication, mutation* and *competition*.

Simulations operate in discrete time intervals *t*, where *t* = 15 minutes and each simulation is run for 100 years. At time interval *t* = 0 the simulator is initialised with an mtDNA_*wild*_ population of copy number *CN*_*wild*_ = 1, 000. In each subsequent interval *t >* 0 mtDNA can replicate (under certain conditions) and randomly mutate. Random deletion mutations lead to heteroplasmy within the organelle such that the overall mtDNA population comprises both mtDNA_*wild*_ and mtDNA_*del*_. Deletion mutations in our simulation are idealised, in that, an mtDNA_*del*_ is half the size of its parent.

### 3.1 Expiration

mtDNA operate in a hostile environment and are subject to oxidative stress due to free radicals. Consequently, mtDNA *expire* with a half-life of 10- 30 days [21, 31]. In our simulation we subject mtDNA to random damage pursuant to a 30 day half-life. mtDNA aging is simulated by assigning a time-to-live (TTL) to each mtDNA which is decremented each time interval *t* according to a Bernoulli trial success:

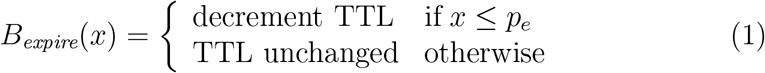

where 0 *≤ x ≤* 1 is a uniformly distributed random number. When the TTL reaches zero, the mtDNA expires. The TTL and the probability *p*_*e*_ are calibrated to yield a half-life of 30 days (see Table 1).

**Table 1:** Simulation parameters, fixed for all simulations.

| Parameter | Value | Comment |
| --- | --- | --- |
| $t_{max}$ | 3,504,000 | Simulation run length (100 years) |
| $p_c$ | 0.01 | Cloning probability $h^{-1}$ |
| $C_{max}$ | 10,000 | Organelle capacity |
| $N_{wild}$ | 1,000 | Initial mtDNA<sub>wild</sub> population |
| $T_{wildRepl}$ | 2 hours | Wild-type replication time ($4 \times t$) |
| $TTL$ | 10 | Equiv. 30 day half-life |
| $p_e$ | 0.0034 | |

### 3.2 Replication

Cells need to maintain a population of mtDNA despite mtDNA having a shorter lifespan than its host cell. mtDNA replicate by cloning, thus, replenishing the population within the organelle. In each time interval *t >* 0 an mtDNA can replicate subject to following conditions:

- The cell has an energy deficit.
- The population of mtDNA does not exceed the organelle’s capacity *C*_*max*_.
- The mtDNA is not currently undergoing cloning.

The cell’s demand for energy is modelled as a consumer/producer mechanism. Each mtDNA_*wild*_ produces *e*_*wild*_ while the cell consumes *E*_*cell*_ every *t*. Thus, for each *t* there is an energy level *E*(*t*), such that:

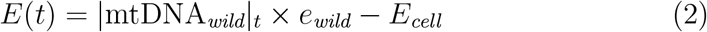

where |*mtDNA*_*wild*_|_*t*_ is the number of mtDNA at time *t*. If energy for the cell is in deficit, cloning is enabled but if there is a surplus cloning is disabled. The energy deficit/surplus mechanism maintains *CN*_*wild*_ for a given *e*_*wild*_ and *E*_*cell*_. The cloning deficit flag *CL*_*def*_ is set accordingly:

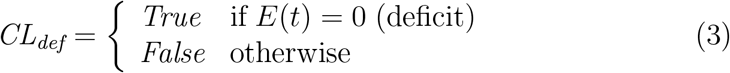

If *C*_*max*_ is the capacity of the organelle, then cloning is enabled only if |*mtDNA*|_*t*_ *≤ C*_*max*_ and is disabled otherwise. Cloning is re-enabled once |*mtDNA*|_*t*_ drops below *C*_*max*_ due to attrition of mtDNA. Thus, the cloning capacity flag *CL*_*cap*_ is set accordingly:

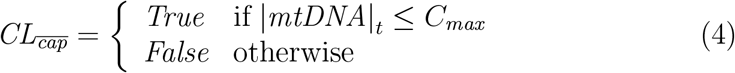

where the 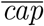 subscript means *not* exceeding capacity.

Cloning is a sequential process, that is, an mtDNA cannot simultaneously clone multiple times and must complete its current cloning process before entering the next. The time to clone is proportional to an mtDNA’s size; given that mtDNA_*del*_ are smaller than mtDNA_*wild*_, deletions have a replicative advantage over wild-type [23]. An mtDNA_*wild*_ is 16,569 nucleotides in size and takes two hours to replicate. Any mutated child is half the size of its parent and, therefore, takes half the time to replicate. For any mtDNA *m*, we compute the replication time (busy period) *T*_*busy*_(*m*) with the expression:

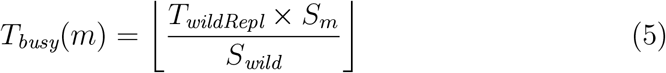

where *S*_*m*_ is the size (in nucleotides) of *m*, S_*wild*_ = 16, 569 is the size of mtDNA_*wild*_ and *T*_*wildRepl*_ = 8 time intervals (two hours). Any mtDNA *m* undergoing cloning enters a *busy* state for *T*_*busy*_(*m*) time intervals. The busy counter for *m* is initialised to *m*_*busy*_ = *T*_*busy*_(*m*) upon entering cloning. *m*_*busy*_ is decremented at each subsequent time interval *t* until it reaches zero.

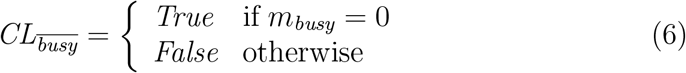

where the 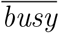 subscript means *not* busy.

*CL*_*def*_ and 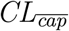 turn cloning off organelle-wide, whereas, *CL*_*busy*_ only effects a specific mtDNA. Cloning is enabled if:

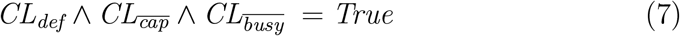

whereby a Bernoulli trial is run for the mtDNA:

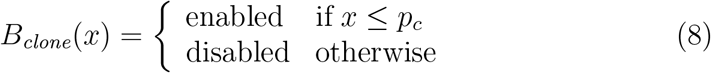

where 0 *≤ x ≤* 1 is a uniformly distributed random number. *p*_*c*_ = 0.01 and is sufficient to counter the attrition rate and maintain a population (when not in competition). A success in *B*_*clone*_(*x*) causes the mtDNA to replicate.

### 3.3 Mutation

As our model is based upon MFRTA, we assume free radical damage causes deletion mutations. This is abstracted in our simulator by a Bernoulli trial with probability *p*_*m*_. Mutation takes place upon the success of the Bernoulli trial:

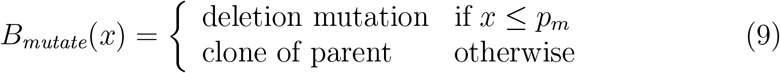

where 0 *≤ x ≤* 1 is a uniformly distributed random number.

When mtDNA undergo cloning, mutation can occur which gives rise to a new species of mtDNA. In this paper we focus solely on mtDNA_*del*_. mtDNA_*del*_ expire and replicate in the same way as mtDNA_*wild*_, thus, mutant species populations can grow in the organelle alongside the mtDNA_*wild*_. mtDNA_*del*_ can also mutate, yielding a larger deletion species.

### 3.4 Fission amd fusion

We implement two separate mitochondrial models, namely, an open mitochondria (OM) and a closed cristae (CC) model. In the OM model mtDNA can freely drift randomly throughout the organelle. With the OM model, mtDNA are stored as a flat *list* and each mtDNA is processed in turn. In each time interval the list is randomly sorted prior to processing in order to mitigate any order bias. This feature of the algorithm also serves to *mix* the mtDNA population throughout the organelle.

For the CC model, the mitochondria is divided up into compartments in order to model closed cristae within an organelle. When the initial population is generated, each mtDNA is randomly assigned to a cristae. mtDNA can replicate/mutate within the confines of their assigned cristae.

Periodically, a cristae may fission momentarily forming two fragments. One fragment remains attached to the main organelle while the other is detached and then *fused* to some other cristae chosen at random. For convenience, each fragment comprises half^1^ the original cristae mtDNA population.

Each mtDNA from the cristae fragment that fissioned, continues to replicate (and mutate), in its newly assigned cristae.

We also implement selective mitophagy whereby we perform a health assessment of a fissioned fragment. A cristae fragment is deemed unhealthy if it contains *no* mtDNA_*wild*_ and healthy, otherwise. A healthy fragment is fused back to the organelle (albeit, to another cristae) but an unhealthy one will undergo mitophagy.

### 3.5 Competition

Enforcing a maximum capacity (*C*_*max*_) in the organelle creates competition for space between species. Initially, there is an abundance of space within the organelle and there is spare capacity for all species of mtDNA. However, as the mtDNA_*del*_ replicate, the aggregate mtDNA population can reach *C*_*max*_. We define the moment when the population reaches *C*_*max*_ for the first time as the *competition point*. At this point, space within the organelle becomes scarce and is only available after some attrition. Each species, therefore, must compete for space when cloning. When the organelle enters competition, selfish proliferation of mtDNA_*del*_ impacts the ability of mtDNA_*wild*_ to maintain energy levels within the cell. The cell dies if the mtDNA_*wild*_ population drops below a threshold (in our simulation, *N*_*wild*_*/*3). This applies to both OM and CC models.

For the CC model, however, each cristae has a capacity limit of *CR*_*max*_ such that *CR*_*max*_ = *C*_*max*_*/N*_*CR*_ and *N*_*CR*_ is the number of cristae in the organelle. In our simulation we set *C*_*max*_ = 10, 000 and *N*_*CR*_ = 1, 000, thus, *CR*_*max*_ = 10. In a cristae, *CR*_*max*_ is enforced in the same way as *C*_*max*_, that is, cloning in the cristae is disabled if the number of mtDNA in the cristae exceeds this limit. Cloning is enabled after some attrition in the cristae.

For convenience, we assume that a fissioned cristae, while initially reduced in size, will grow back to it’s original size *CR*_*max*_. Conversely, cristae that fuses with a fissioned fragment will reduce down to *CR*_*max*_. While the number of mtDNA in the cristae may exceed *CR*_*max*_ after fusion, cloning is disabled in the cristae until attrition reduces the population below the capacity of the cristae.

### 3.6 Parameters

This subsection summarises the parameters used in the simulation. Table 1 shows the parameter values common to both OM and CC models, whereas, Table 2 shows the parameters specific to the CC model. The mutation probabilities *p*_*m*_ for low, medium and high dementia predisposition levels is shown in Table 3.

**Table 2:**
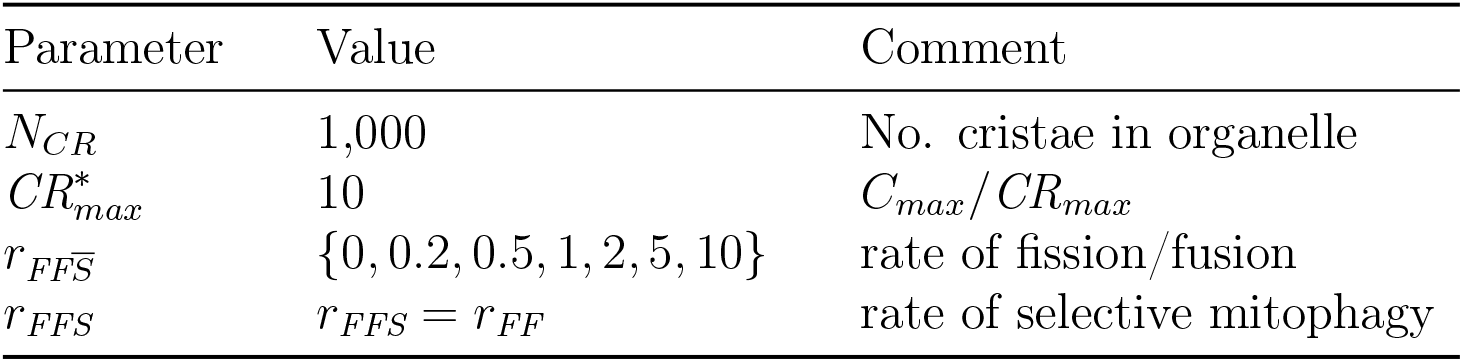
Parameters for CC model.

| Parameter | Value | Comment |
| --- | --- | --- |
| $N_{CR}$ | 1,000 | No. cristae in organelle |
| $CR_{max}^*$ | 10 | $C_{max}/CR_{max}$ |
| $r_{FF\bar{S}}$ | $\{0, 0.2, 0.5, 1, 2, 5, 10\}$ | rate of fission/fusion |
| $r_{FFS}$ | $r_{FFS} = r_{FF}$ | rate of selective mitophagy |

**Table 3:** Mutation probabilities associated with predisposition to dementia. For most of the experiments we run, we only consider low, medium and high predispositions. v/high (very high) is only used simulate selective mitophagy in an enviroment of high stress.

| Predisposition | $p_m$<br>($h^{-1}$) |
| --- | --- |
| low | $1 \times 10^{-4}$ |
| medium | $1.5 \times 10^{-4}$ |
| high | $2 \times 10^{-4}$ |
| v/high | $1 \times 10^{-2}$ |

We run 200 replications of the simulation for each factor and level. Source code is available in the Git repository: `git clone` https://agholt@bitbucket.org/agholt/mitosim2022.git.

There is little consensus in the literature as to what cell loss rates, both global and local, constitute dementia [18, 15, 59, 3]. We use neuron loss ranges from Arendt *et al*. [4] and Holt *et al*. [24], that is, 20-40% loss, where the lower limit is the onset of dementia and the upper is severe. Below 20% is healthy aging and beyond 40% it is unlikely the host itself survived, however, we run the simulation for the full 100 years, regardless.

## 4 Analysis

We present the results of our simulations for a CC mitochondrion that undergoes fission and fusion, and compare it to an OM model. In our first set of simulations we focus on 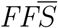 and *FFS* in the second set.

The graphs in Fig. 1 show the experimental cummulative distribution function (ECDF) for neuron lifespan. These graphs give us the rate of neuron loss for predisposition and 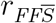. The black lines show the neuron loss for the OM model and the coloured lines represent the neuron loss for various values of 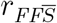.

**Figure 1:**
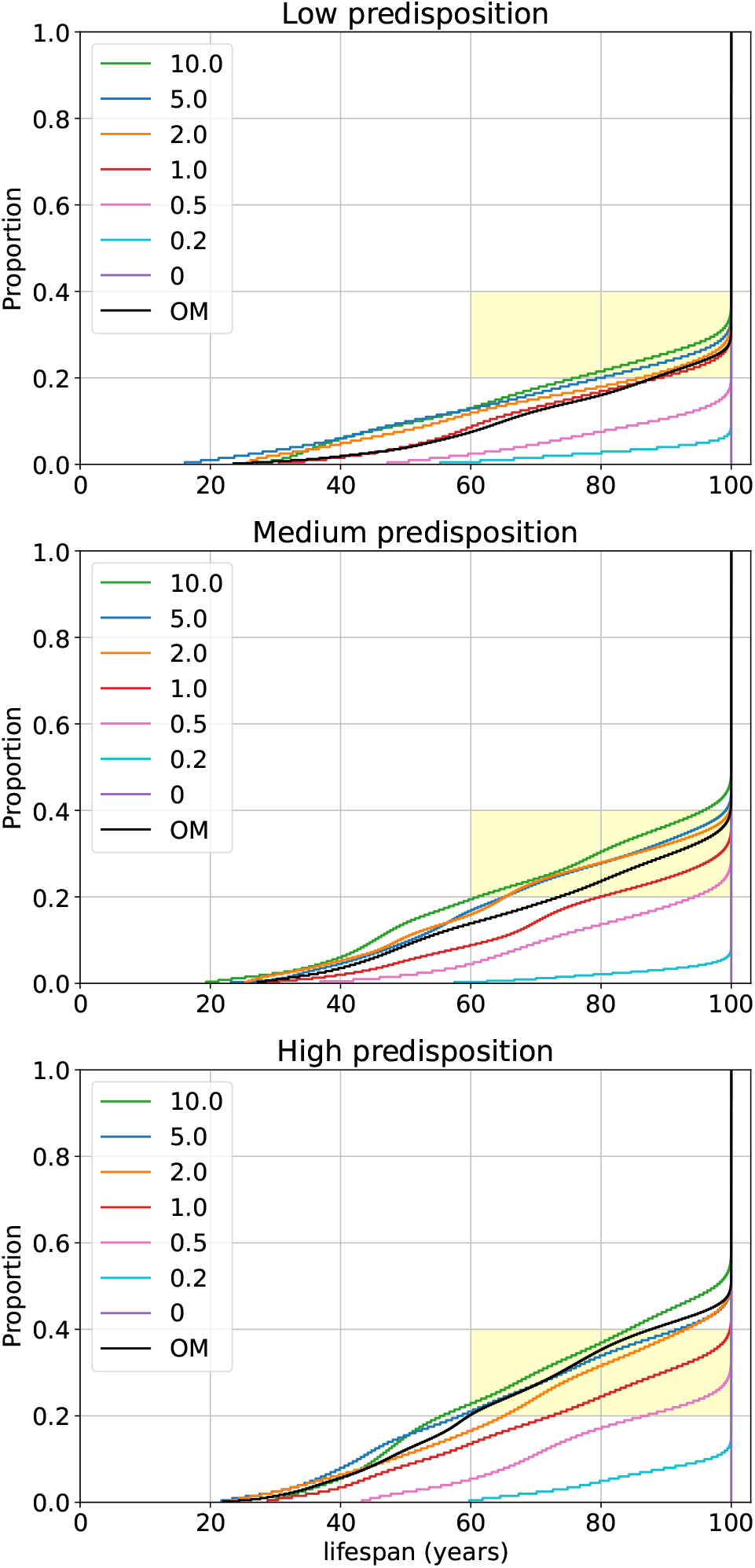
Neuron lifespan for low, medium and high predisposition. Black lines are OM and coloured lines are CC for varying 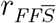. The yellow shaded area represents the region of the system where the cognitive decline in the host would be classed as dementia for the elderly.

Neuron loss and, thus, the prevalence of dementia, increases with predisposition and is a result we have verified in previous papers [25]. For CC, neuron loss increases with 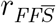. For 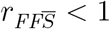, there is little cognitive decline with age and, only in the worst case, onset of dementia is in extreme old age. Indeed, for 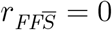 (no mixing) there is no neuron loss.

The severity of dementia, and its onset, is brought forward with higher mixing rates, albeit, neuron losses appear to reach a limit for 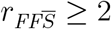. The losses for high mixing rate 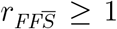 in CC are comparable to OM, albeit, the value of 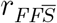 where CC and OM are comparable increases with predisposition. There is clearly a difference in the mtDNA proliferation dynamics between mixing by 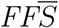 in CC and mixing by diffusion in OM.

The graphs in Fig. 2 show heteroplasmy for deceased cells measured as the total number of mutant species throughout the lifespan of the host. We confine our analysis to deceased cells because survived cells have, typically, low mutation levels and thus low heteroplasmy. In both CC and OM, heteroplasmy increases with predisposition; again this is a result consistent with previous research (cited above).

**Figure 2:**
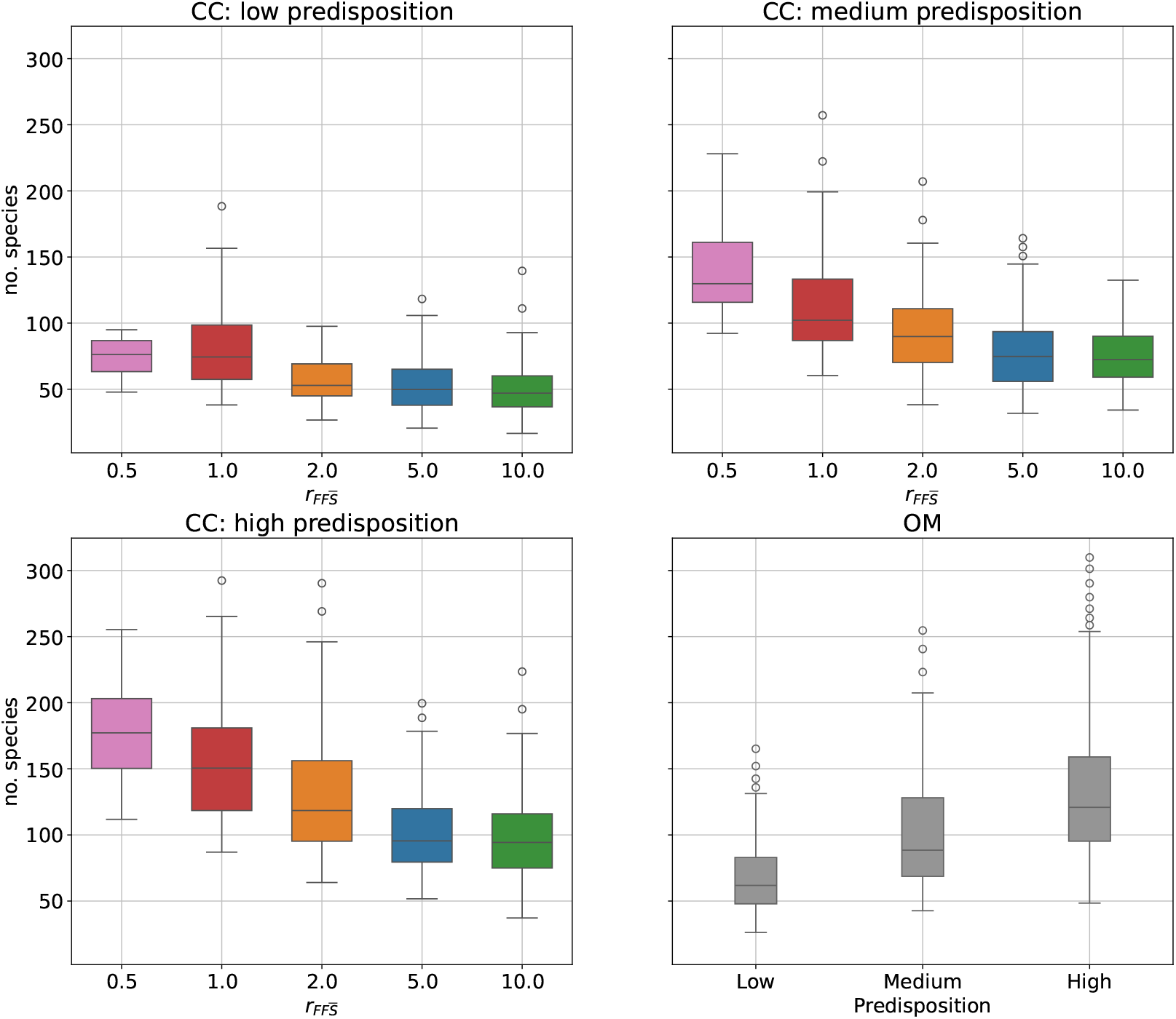
Heteroplasmy. Top left: Low predisposition CC. Top right: Medium predisposition CC. Bottom left: High predisposition CC. Bottom right: All predispositions OM.

The important result here is that heteroplasmy declines with 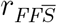. Deceased neurons will have similar levels of mutants regardless of predisposition and values of 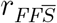; heteroplasmy, therefore, gives us an indication of clonal expansion. From Nido *et al*. [42] we expect 1-4 clonal expansions with a background of heteroplasmy and this is what we observe in our simulation results. The lower the heteroplasmy, the more the mutant population is dominated by larger, but fewer, clonal expansions. The converse is true of high heteroplasmy. Thus, higher values of 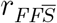 result in lower heteroplasmy, albeit there appears to be a limit at 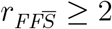. For each level of predisposition, the level of heteroplasmy of CC for 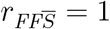 is comparable with OM.

Here we present the *FFS* results. We focus on a host with high predisposition (*p*_*m*_ = 2 *×* 10^*−*4^) and a subset of values of 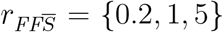 where 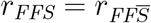.

The graph in Fig. 3 shows the *FFS* results along side the results for 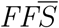 (extracted from the graphs in Fig. 1). For all rates of *r*_*FFS*_ = {0.2, 1, 5}, *FFS* resulted in no neuron loss. Our *in silico* selective mitophagy function is effective in maintaining healthy mitochondria. However, given the prevalence of dementia observed in the human population, it appears to be somewhat *too* effective.

**Figure 3:**
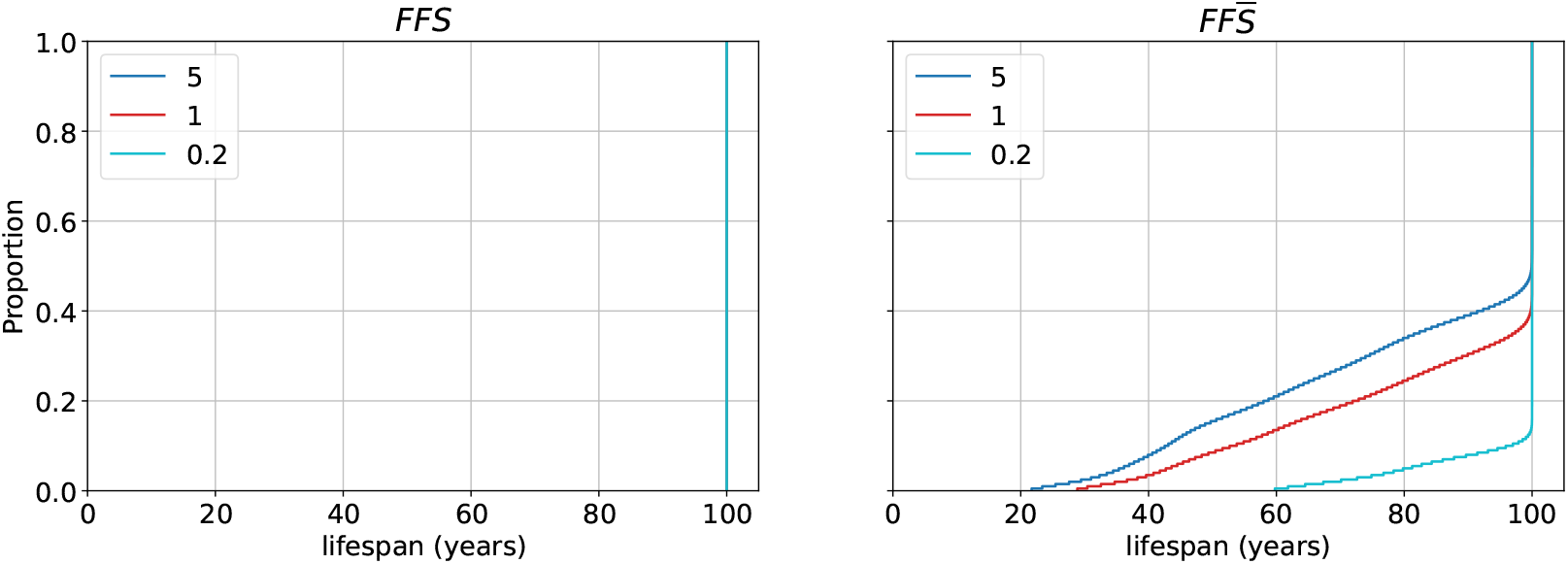
Lifespan *FFS* compared to 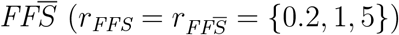

We considered the possibility that we had underestimated the *hostility* of the environment mtDNA operate in. Thus, we ran a set simulations for a *very high* predisposition (v/high) with a mutation rate *p*_*m*_ = 10^*−*2^ *h*^*−*1^ and 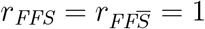^2^.

However, even at this level of predisposition, we only had one neuron loss in 200 simulation runs. When we analysed the heteroplasmy for v/high predisposition, we saw that the number of mutant species was an order of magnitude higher than low/medium/high predisposition for comparable values of 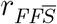 (see Fig. 4). For the one simulation run that resulted in neuron loss, there was one predominant clonal expansion. A snapshot taken at the end of the simulation run, showed that the background heteroplasmy accounted for 60% of the mtDNA population while the clonal expansion accounted for only 37% (the remaining 3% being the wild-type). This does not fit with the empirical evidence of low levels of background heteroplasmy dominated by just a few clonal expansions [42].

**Figure 4:**
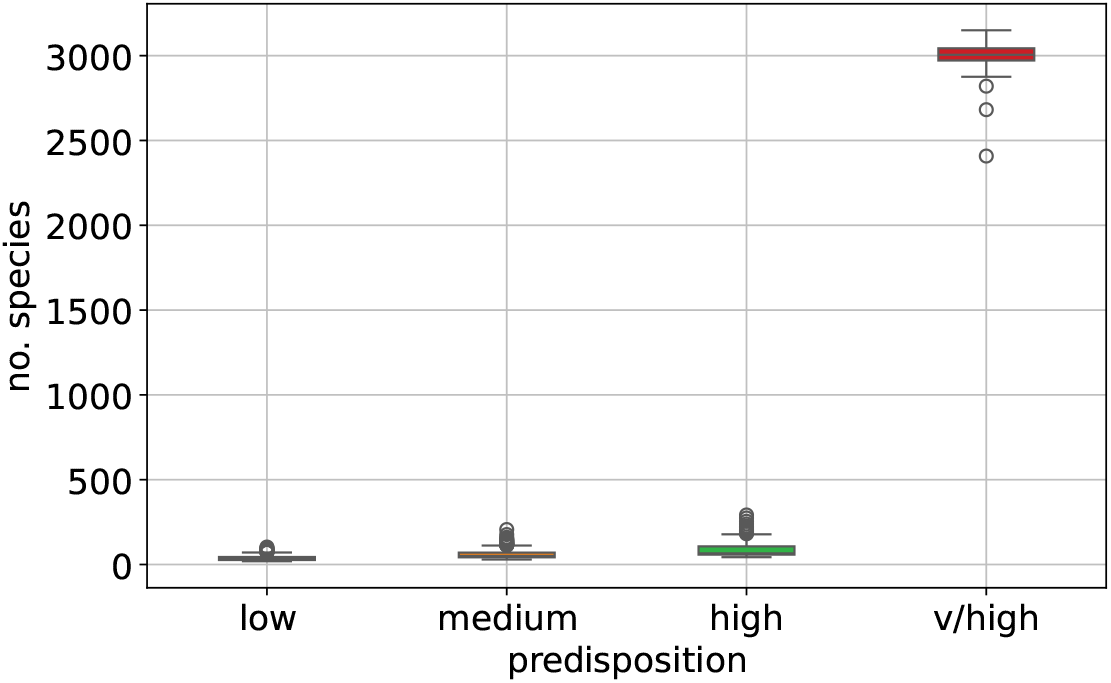
*FFS* heteroplasmy of very high (v/high) predisposition (*p*_*m*_ = 10^*−*2^ *h*^*−*1^) and FF rate of 1 *t*^*−*1^, survived and deceased cells combined. Compared to FF with no selective mitophagy (low/medium/high predisposition, *r*_*FF*_ = 1 *t*^*−*1^, survived and deceased cells).

Neuron loss necessitates clonal expansion and our selective mitophagy function was effective at preventing the expansion of a mutant species. What appeared to make *FFS* so effective in our simulation was high rates of selective mitophagy *r*_*FFS*_. In the simulations above, 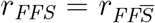. High levels of neuron loss attributed to high 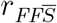 in 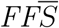 are entirely mitigated by high *r*_*FFS*_ in *FFS*. Low 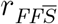 in 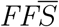 yields low neuron loss making *FFS* largely redundant even at low *r*_*FFS*_.

Thus, in our final experiment we disassociated *r*_*FFS*_ from 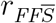 by reducing *r*_*FFS*_ to 1*/*1000^*th*^, 1*/*100^*th*^ and 1*/*20^*th*^ of 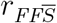, where 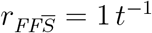. For approximately five selective mitophagy events a day (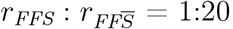) there was no neuron loss. At (approximately) one a day (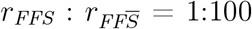), there was a small amount of loss in old age. At only one event every 10 days, the levels of neuron loss are similar to 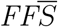. However, all this means is that, no selective mitophagy yields similar results to *almost* no selective mitophagy.

## 5 Discussion

We investigated the effect of fission and fusion in mitochondria on long term neuron loss in humans. We modelled mitochondria as set of closed cristae (CC). Unlike an open mitochondria (OM), where mtDNA are free to drift randomly throughout the organelle, the CC model confines mtDNA to cristae. Organelle-wide mixing of mtDNA is only possible through the process of fission and fusion.

We simulated the proliferation of mtDNA_*del*_ for various predispositions (low, medium, high) and for various rates of mixing and selective mitophagy (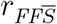 and *r*_*FFS*_, respectively). We compared the neuron loss results for 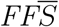 with 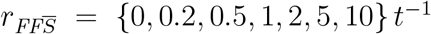 with the OM model. The results showed that the OM model yields roughly the same levels of neuron loss as the 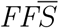 model at high mixing rates, that is, 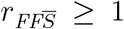. Surprisingly, low rates 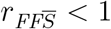 reduced neuron loss such that the onset of dementia was delayed until late old age and severity was significantly reduced. Indeed, for the no mixing case 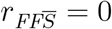, there was no neuron loss at all.

That the prevalence of dementia increases with 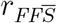, seemed counter intuitive at first. However, there is precedence for this result. Observations *in vitro* show that mitochondria, when quiescent, tend to take the form of an integrated threadlike network [7] until subject to stress, resulting in a decrease in the ΔΨ and causing the mitochondrion to fragment [46, 57].

Many compounds can induce stress in a mitochondrion by decreasing the ΔΨ, causing it to fragment [46, 57]. Among farm workers, rotenone exposure is associated with a Parkinson’s-like neurodegenerative pathology [52, 56]. Furthermore, rotenone is also used to model Parkinson’s disease in animals [10, 13] and is often used as a positive control for depolarization and fragmentation of the mitochondrion in screening assays [26, 46]. These observations, along with many others [65, 17, 44, 58], suggest a link between compounds that cause mitochondrial fragmentation *in vitro* and neurodegenerative pathologies *in vivo*.

This raises the question as to the purpose of fission and fusion mixing. In terms of deletion mutations, 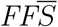 does not maintain mitochondrial function, in fact it does the opposite. Mixing in 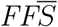 facilitates genetic diversity of the mutant population transferring deletions with a replicative advantage into the otherwise homoplasmic cristae of wild-type.

It is conceivable that 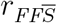 in humans is nominally low *in vivo*. Compounds that increase 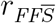 *in vitro* could also do so *in vivo* and could explain why certain medications, taken over many years, are associated with an increase in the prevalence of dementia. Furthermore, those hosts predisposed to higher 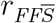 will suffer a higher predisposition to dementia (on top of increased mutation probability).

We ran experiments where mtDNA mixing worked in conjunction with selective mitophagy (*FFS*). Our results showed that *FFS* successfully prevents clonal expansion. However, *in silico FFS* was *too* effective, in that, it eliminated clonal expansion altogether. All neurons survived to the end of the simulation, regardless of predisposition and 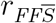. Thus, the prevalence of dementia that we observe in the general population should not be so high.

We attribute the success of *in silico FFS* to high levels of *r*_*FFS*_ given that we set 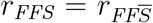 in our initial *FFS* experiments. When we slowed down *r*_*FFS*_ relative to 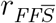, neuron loss resulted. However, we had to slow *r*_*FFS*_ down considerably relative to 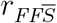 in order to see levels of cognitive decline similar to the degrees of dementia observed in human populations.

We speculate that the ratio 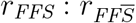 may differ from host to host and may contribute (along with mutation probability) to predisposition. However, given the results in Fig. 5, this would necessitate some amount of fine tuning.

**Figure 5:**
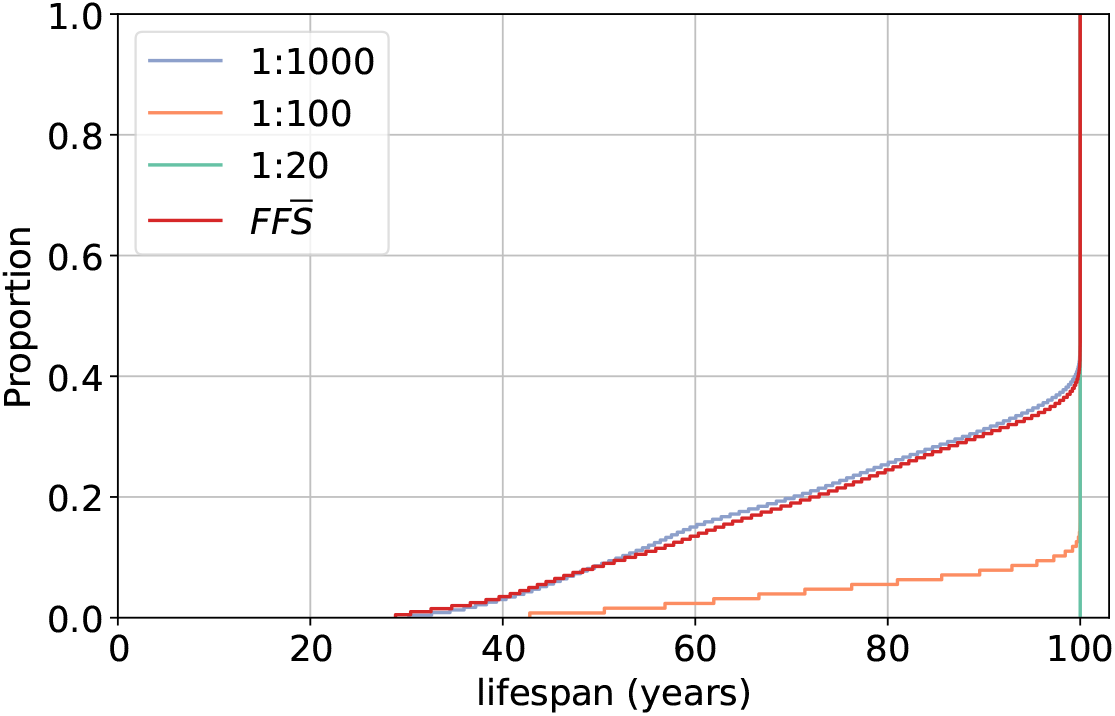
Neuron lifespan 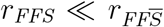. Graphs show results 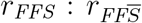 rate ratios of 1:1000, 1:100 and 1:20, where 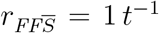. For comparison, the 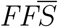 case is included (for 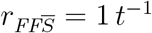).

The initial mtDNA population in a host will be homoplasmic having passed through the germline bottleneck [35, 64, 47, 9]. There will, therefore, be little or no phenotypic variation in mitochondria fragments in young hosts given that there would not have been a significant accumulation of mutations. Indeed, we can see from Fig 1 that neuron loss is low (indicative of low clonal expansion) below reproductive age. Thus *FFS* is redundant in the young and the level of *r*_*FFS*_ is inconsequential.

Viewed through the lens of natural selection, if *FFS* is a dementia mitigation mechanism then it would have to be a functional adaptation brought about by some selection pressure. However, dementia is a disease of the elderly and the force of natural selection declines with age [62, 22]. Thus, it is difficult to conceive of a functional adaptation that does little for the young and declines in effectiveness in the elderly.

There is no clear concensus as to why mitochondria undergo fission and fusion, however, it has been demonstrated that fusion is dependent on ΔΨ and that, fragments with a reduced ΔΨ, cannot undergo fusion, probably because of depleted ATP [54]. While our results indicate that the process of fission and fusion is not likely to be for the purpose of removal of mtDNA_*del*_ by selective mitophagy, or even to facilitate complementation, fission and fusion is still vital as its failure can be fatal [6].

A possible function of fission and fusion is to facilitate cellular adaptation to changing energy demands and other metabolic needs [61, 36] though it is not clear why this would be the case. The most extreme mitochondrial fragmentation is seen *in vitro* in response to cell stress [19, 63], but again, it is not clear why fragmentation is an appropriate response to stressors.

## 6 Conclusions

In this paper we modeled a mitochondria as collection of closed cristae (CC). In contrast to an open mitochondria (OM) model where mtDNA can diffuse freely throughout the organelle, the cristae keep mtDNA confined. mtDNA can only relocate to another part of the organelle (cristae) by means of fission and fusion. We compare the proliferation of mtDNA_*del*_ within the CC model (without selective mitophagy) for various 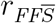 with that of the OM model. In each case we record the neuron loss due to the proliferation of mtDNA_*del*_ over the course of a human lifetime (100 years).

Neuron loss increases with 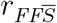 showing that, CC and OM yield comparable levels of neuron loss at high mixing rates (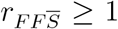). At low rates (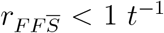), neuron loss is almost non-existent. This result may explain why certain pharmaceuticals and pesticides are associated with increased risk of dementia. Such compounds place stress on the mitochondrion causing excessive fragmentation.

The fusion process that follows facilitates the distribution of damaged mtDNA throughout the organelle which, in turn, promotes clonal expansion.

If we introduce selective mitophagy to the CC model, neuron loss is completely eliminated. Selective mitophagy prevents clonal expansion, which is a requisite of neuron loss due to the proliferation of mtDNA_*del*_. Therefore, *FFS* does not predict the prevalence of dementia observed in human populations, whereas, OM and 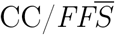 with high 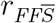 do.

It is possible that our simulation of *FFS* is overly idealised and *in vivo FFS* is less effective and merely slows the decline of cognitive dysfunction. We simulated this by decoupling 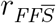 and *r*_*FFS*_ such that *r*_*FFS*_ was slower than 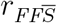. However, we had to reduce *r*_*FFS*_ considerably in order to produce any appreciable level of neuron loss. Setting the *r*_*FFS*_ such that it would yield levels of neuron loss associated with healthy ageing while avoiding dementia in old age would require a significant amount of fine tuning. If this were the case, then selective mitophagy must fail in significant parts of the elderly population given the prevalence of dementia in that demographic. Thus, our *in silico* selective mitophagy function operates on an *all or nothing* basis. This leads us to question the conventional wisdom that fission and fusion acts as a purification mechanism to remove defective mtDNA. From an evolutionary perspective, selective mitophagy is hard to explain. A functional adaptation that is unnecessary in the young and suboptimal in post-reproductive age, would struggle to yield any selective advantage.

Fission and fusion may perform a necessary biological function but we doubt it is a counter measure for dementia in the elderly given that the mixing feature appears to facilitate its prevalence. While selective mitophagy counters the effect of the mixing process the results *in silico* are not representative of the reality *in vivo*.

Our results suggest that dementia is a byproduct of some form of antagonistic pleiotropy, whereby, genes can confer benefits in early life but have deleterious effects in old age. Dementia, therefore, is a consequence of the trade-offs inherent in the evolutionary processes that shapes human development and aging.

## Footnotes

1 If there is an odd number of mtDNA then the attached fragment comprises the larger amount.

2 We appreciate, that if we have underestimated the hostile environment, then the level of predisposition for which *p*_*m*_ = 10^*−*2^ would be considered *normal*. However, we maintain the nomenclature low to v/high for convenience.

